# PSC-RED, an Albumin-Free Robust Erythroid Differentiation Method to Produce Enucleated Red Blood Cells from Human Pluripotent Stem Cells

**DOI:** 10.1101/616748

**Authors:** Emmanuel N Olivier, Shouping Zhang, Zi Yan, Sandra Suzuka, Karl Roberts, Kai Wang, Eric E Bouhassira

## Abstract

Cultured red blood cells (cRBCs) have many potential applications in transfusion medicine and drug delivery. We report that we have developed chemically defined, albumin-free Robust Erythroid Differentiation (RED) methods to produce enucleated cRBCs from human induced pluripotent stem cells (iPSCs). Human iPSC-derived cRBCs produced with either the short or long variation of the RED protocol respectively express embryonic/fetal or a mixture of fetal and adult hemoglobins. The long version of the protocol produces up to 50% of enucleated cells at an unprecedented yield. RED is scalable and relies on inexpensive components and therefore dramatically increases the feasibility and economic viability of all translational applications of cRBCs.

**Highlights:**

- PSC-RED: A chemically-defined, albumin-free Robust Erythroid Differentiation (RED) methods to produce cRBCs from human induced pluripotent stem cells.
- PSC-RED produces up to 50% enucleated cells at an unprecedented yield.
- PSC-RED is scalable and relies on inexpensive components and therefore increases the feasibility and economic viability of translational applications of cRBCs.

## Introduction

Cultured red blood cells (cRBCs) are useful to study erythroid disease mechanisms, hold great promise as reagent cells to diagnose allo-immunization and act as a potential source of invaluable cells carrying rare blood groups that are necessary to transfuse allo-immunized patients (Bouhassira, 2012; Zeuner et al., 2012). More general transfusion applications might also become possible in the future but the costs and technical difficulties involved in producing enough cells to treat large number of patients are major issues (Timmins and Nielsen, 2009). Genetically modified cRBCs that express therapeutic proteins are another highly promising avenue of research, because drugs carried by RBCs are spatially restricted to the lumen of the cardio-vascular system, are shielded from the immune system, and can have a longer half-life than drugs delivered directly to the plasma (Bouhassira, 2012; Magnani, 2017; Rousseau et al., 2014). Importantly, relatively small numbers of such cells could be clinically useful, greatly decreasing the technical barriers to translation.

To be of translational value, methods to produce cRBCs must yield enucleated cells that are as similar as possible to cells produced *in vivo*. Human cRBCs can be produced by expansion of the hematopoietic stem and progenitor cells present in primary cord blood and adult peripheral blood mononuclear cells (MNCs) (Carotta et al., 2004; Dolznig et al., 2005; Giarratana et al., 2005; Migliaccio et al., 2010; Miharada et al., 2006) or from immortal cells. MNCs are not immortal and would therefore need to be constantly collected from multiple donors to ensure a permanent supply of cells if they were used to produce cRBCs at an industrial scale. The use of multiple donors would complicate the production of cRBCs because the genetic variability of the donors would affect the yield, and would introduce unnecessary heterogeneity in the cRBCs. It would also increase the cost and carry a risk of contamination by unknown or emerging pathogens that cannot be completely eliminated even with stringent donor screening. We and others have therefore focused on producing cRBCs from more permanent cell sources such as embryonic stem cells (Kaufman et al., 2001; Qiu et al., 2005; Zambidis et al., 2005), induced Pluripotent Stem Cells (iPSCs) (Fujita et al., 2016; Huang et al., 2014; Olivier et al., 2016; Qiu et al., 2008; Rousseau et al., 2014), self-renewing (England et al., 2011) or immortalized progenitors (Hirose et al., 2013; Kurita et al., 2013; Trakarnsanga et al., 2017).

Human iPSCs are an attractive source of cells for cRBCs production, because they are immortal, can be reproducibly generated from any donor, and because they can be cultured by well-understood methods in chemically defined conditions (Chen et al., 2011). In addition, they are karyotypically stable (Martin, 2017) and genetically modifiable. However, complex differentiation procedures relying on undefined animal or human components, low rates of *in vitro* enucleation (<5%), and an immature globin expression phenotype (Qiu et al., 2008) have hampered the translational use of iPSC-derived cRBCs.

To decrease costs, culture methods must be simple and reproducible, which is best achieved by using chemically defined components. We report here that we have developed scalable chemically defined, albumin-free Robust Erythroid Differentiation (RED) protocols to differentiate human iPSCs into large numbers of enucleated RBCs.

## Materials and methods

### Samples

Peripheral blood (PB) cells were obtained from healthy donors under a protocol approved by the Albert Einstein College of Medicine IRB.

Mononuclear cells from whole blood were purified by histopaque (Sigma-Aldrich) according to the manufacturer instruction, and residual RBCs were lysed by addition of 5 volumes of cold (4°C) RBC lysing buffer (NaHCO3 790mg/L and NH4Cl, 7.7g/L) for 10 minutes. Aliquots containing about three million cells were then frozen and stored in liquid nitrogen.

### Reagents

The suppliers for all reagents are provided in Table S2.

### Culture conditions

All cultures were performed at 37C, in 5 or 10% CO_2_ depending on the presence of stable glutamine in the medium. Since our original protocol was performed in 5% oxygen, all experiments described above were also performed in 5% oxygen, in accordance with previously published results (Vlaski et al., 2009).

### Induced-pluripotent stem cells

iPSCs were reprogrammed from peripheral blood mononuclear cells using the Sendai virus approach (CytoTune-iPS 2.0 Sendai Reprogramming Kit - Thermo Fisher Scientific) according to the manufacturer instruction. 5 lines of iPSCs (NY22, OM1, OM2, OM3 and OM4) and three sub-lines of OM1, all generated from healthy controls, were used during these experiments. The vast majority of experiments were performed with lines NY22 and OM1. All lines enucleated at a high rate although we noted differences that might have been associated with different growth rates between lines.

### Pluripotent stem cell culture

human pluripotent stem cells (hPSCs) were maintained undifferentiated in E8 medium (Chen et al., 2011) on Vitronectin (life technologies) and passaged using EDTA every 3-4 days depending on their confluence stage.

## Differentiation of iPSCs into erythroid cells: Short PSC-RED protocol

**On day -1**: Three-day-old hPSC colonies were dissociated with 5mM EDTA in PBS for 6 minutes. The EDTA was then removed and replaced with 5 mL of E8 medium and the well was thoroughly flushed with a 5 mL serological pipet by pipetting up and down 10 times. Small clumps were generated to produce small colonies of about 50 cells on day 0. The cells were then plated at 1-2×10^5^ cells/well in 2mL/well of E8 medium on vitronectin in tissue culture treated six-well plates (Falcon), which are used throughout the protocol. After plating, the cells were allowed to attach overnight.

**On day 0**: Differentiation was induced by replacing the E8 medium with IMIT medium, containing **supplement 1** (Bone Morphogenic Protein 4 (BMP4) (10ng/mL), Vascular Endothelium Growth Factor 165 (VEGF) (10ng/mL), basic Fibroblast Growth Factor (bFGF) (10ng/mL), Wnt3A (5ng/mL), Wnt5A (5ng/mL), Activin A (5ng/mL) and GSK3β Inhibitor VIII (2μM) (Olivier et al., 2016))

Before inducing the differentiation, the culture was inspected to ascertain that most of the colonies contained about 50 cells or less. One well of the culture was sacrificed for cell counting in order to calculate the yield of cells at the end of the experiments.

**On day 2**, 6x concentrated **supplement 2** in IMIT was added to each well to bring the final concentration of fresh cytokines to 20ng/mL of BMP-4, 30ng/mL of VEGF, 5ng/mL of Wnt3A, 5 ng/mL of Wnt5A, 5ng/mL of Activin A, 2μM of GSK3β Inhibitor VIII, 10ng/mL of bFGF, 20ng/mL of SCF and 0.4ng/mL of β-Estradiol.

**On day 3**, the cells were dissociated with TrypleSelect 1× for 5-10 minutes at 37C. After the addition of 10 mL of PBS, cells were centrifuged for 3 minutes at 250g, the supernatant was discarded and the cells re-suspended in fresh IMIT medium containing **supplement 3** (BMP4 (20ng/mL), VEGF (30ng/mL), bFGF (20ng/mL), SCF (30ng/mL), Insulin-like Growth Factor 2 (IGF2) (10ng/mL), Thrombopoietin (TPO) (10ng/mL), SB431542 (3μM), Heparin (5μg/mL), IBMX (50μM) and β-Estradiol (0.4ng/mL) and plated at 1.10^5^ cells/mL of a tissue culture treated six wells plate (3 mL per well).

**On day 6**, the cells were centrifuged for 3 minutes at 350g and re-suspended at 5.10^5^/mL in fresh IMIT medium containing supplement 3 without SB431542 but with 30nM of UM171.

Between day 6 and day 10, the cells were diluted to 0.5.10^6^/mL any time they reached more than 1.5.10^6^cells/mL by addition of the same medium and supplement. An additional dose of S3 supplement (provided from a 6x concentrated stock) was added at day 8 to fully renew the cytokines and small molecules.

**On day 10**, the cells were centrifuged for 3 minutes at 250g, plated at 0.66.10^5^ cells/mL in IMIT containing the SED supplement (SCF 100ng/mL, Erythropoietin 4U/mL, IBMX 50μM and Dexamethasone 1μM). From day 10 to day 17, the cells were diluted to 0.5.10^6^/mL any time they reached more than 1.5×10^6^cells/mL by addition of the same medium and supplement. In addition, 6x concentrated SED supplement in IMIT was added every 2 days to fully renew the cytokines and small molecules.

**On day 17**, the cells were centrifuged for 3 minutes at 250g and plated at a density of about 2×10^5^/mL of IMIT containing the SER supplement (SCF (50ng/mL), EPO (4U/mL) and RU486 (1μM)). From day 17 to 24, the cells were diluted to 0.5×10^6^/mL any time they reached more than 1.5×10^6^cells/mL by addition of the same medium and supplement. In addition, 6x concentrated SER supplement in IMIT was added every 2 days to fully renew the cytokines and small molecules.

**On day 24**, the cells were centrifuged for 3 minutes at 250g and plated at 2×10^5^/mL in R5 medium with the SER2 supplement (SCF (10ng/mL), EPO (4U/mL) and RU486 (1μM)). From day 24 to 31, the cells were diluted to 0.5×10^6^/mL any time they reached more than 1.5×10^6^cells/mL by addition of the same medium and supplement. In addition, 6x concentrated SER2 supplement in R6 was added every 2 days to fully renew the cytokines and small molecules.

**On day 31**, the cells were centrifuged for 3 minutes at 250g and maintained in R5 or R6 medium alone for up to 8 days.

## Long Differentiation protocol

This long protocol is identical to the short protocol but an additional HPC expansion step in added after day 10. This step consists of centrifuging the cells at 250g for three minutes and replating the day 10 cells in IMIT at 2×10^5^/mL in the presence of **supplement 4** (bFGF (5ng/mL), SCF (15ng/mL), VEGF (5ng/mL), TPO (10ng/mL), IGF2 (10ng/mL), Platelet Derived Growth Factor (PDGF) (5ng/mL), Angiopoietin-like 5 (ANGPTL5) (5ng/mL), Chemokine Ligand 28 (CCL28) (5ng/mL), IBMX 30μM, Heparin (5ug/mL) and UM171 (30nM) for one or two weeks. As above, the concentration of cells is kept below 1.5×10^6^cells/mL at all times and cytokines are refreshed every two days by adding 6x concentrated supplement. Cells kept for two weeks in these conditions, were centrifuged and transferred to fresh plates after 7 days to eliminate any attached cells.

After this additional step, the differentiation resumes according to the short protocol day 10. A one-time addition of Granulocyte-Macrophage Colony Stimulating Factor (G-MCSF) (20ng/mL) and Granulocyte Stimulating Factor (G-CSF) (20ng/mL) is, however, necessary to induce maximal proliferation of the HPCs in the SED supplements.

## Analysis and characterization

### Cell enumeration

Cells were counted with a Luna-FL dual channel Automated Cell Counter (Logos) using acridine orange to visualize the live cells and propidium iodide to exclude the dead cells

### Flow cytometry

iPSCs undergoing differentiation were evaluated by FACS using antibodies against CD34, CD36, CD43, CD45, CD71 and CD235a also known as glycophorin A (BD Biosciences and eBioscience).

### Enucleation

The enucleation rate was measured using the DRAQ5 DNA nuclear stain (ThermoFisher) after exclusion of dead cells with Propidium Iodide. The cells were analyzed with a BD FACS Calibur flow cytometer (BD Biosciences) or a DPX10 (Cytek) flow cytometer and the flow cytometry data were analyzed with the Flowjo software.

### Giemsa staining

Erythroid differentiation and enucleation were also assessed microscopically by Rapid Romanovsky staining of cytospin preparations. Cell sizes were estimated on a Nikon TE-2000S microscope using software provided by the manufacturer.

### RBC filtration

To eliminate the nuclei at the end of the experiments, RBCs were filtered using PAL Acrodisc 25mm WBC filters as recommended by the manufacturer. Filtered cRBCs were stored for up to one month with little signs of hemolysis in Alsever’s solution (Sigma).

### HPLC analysis

Cells were washed twice with PBS, and lysed in water by 3 rapid freeze-thaw cycles in dry-ice and a 37C water bath. Debris was eliminated by centrifugation at 16 000g and the lysates stored at −80°C. HPLC were performed as described (Fabry et al., 2003). Briefly, a few μL of lysate containing about 50μg of protein in about 100 μL of 40% acetonitrile and 0.18% TFA was filtered and loaded on a VYDAC C4 column. The globins were then eluted with increasing concentration of acetonitrile during a period of about 80 minutes. The starting elution buffer was programmed to be 80% buffer A and 20% buffer B and to rise to 50% buffer B in 50 minutes. Buffer A = 36% acetonitrile and 0.18% TFA and buffer B = 56% acetonitrile and 0.18% TFA. Globin chain elution was monitored by measuring O.D. at 220 nm.

## Results

### Production of enucleated cRBCs from iPSCs

Olivier et al. (2016) previously published an efficient method to produce cRBCs from iPSCs that relied on commercial medium StemLine II to differentiate iPSCs into successive Hematopoietic Progenitor Cells (HPC) and erythroblasts, and IBIT, a serum-free IMDM-based media containing human-albumin, insulin and transferrin, to terminally differentiate the erythroblasts into cRBCs. However, the use of animal-derived components increases costs and decreases the reliability of the method. Indeed, because of extreme lot variability, we replaced Stemline II during the first steps of the protocol with IBIT, and were able to achieve slightly higher yields of cRBCs. However, this did not solve the problem of reproducibility because lots of human or bovine albumin also proved extremely variable and recombinant albumin can be hard to source and very expensive. We therefore decided to develop a chemically defined protocol that would not require any albumin or other animal components and revisited several steps of the differentiation protocol.

### Elimination of embryoid body formation

To eliminate all animal components from the protocol, we started with iPSCs grown in chemically-defined conditions, instead of iPSCs grown in Stempro hESC, an albumin containing medium, as described previously (Olivier et al., 2016). Because embryoid bodies (EB) do not form readily when iPSCs are grown in chemically-defined conditions (Lin and Chen, 2008), we developed a two dimensional mesoderm induction procedure instead. This first step relies on 0.5mM EDTA in PBS to dissociate iPSCs grown on E8/vitronectin into small clumps, and on plating these small clumps directly into mesodermal differentiation conditions that involve three successive cytokine supplements termed S1 to S3 (Table S1). The size of the iPSC clumps at this stage is important in order to maximize the contact of the differentiation factors with the cells while avoiding the protective colony effect on the cells in their centers. Figure S1 illustrates the morphology of the cells during the first days of the differentiation.

### Elimination of albumin

We initially tested the R5 and R6 media that are described in the accompanying manuscript (Zhang S. et al) and that were designed for the terminal differentiation of cultured erythroblasts, but these media were too poor to support iPSC differentiation into HPCs. We therefore devised IMIT, a medium similar to previously published IBIT but which contains Trolox, an anti-oxidant, and methyl-β-cyclodextrin (Pei et al., 2017) instead of albumin. This last compound helps solubilize lipid soluble molecules and can therefore partly substitute for albumin. Ethanolamine, an important membrane component, was also added because it has been shown to improve cellular growth in serum free media developed for hybridoma culture (Kano-Sueoka, Ta et al., 2001; Murakami et al., 1982; Patel and Witt, 2017).

Comparison of IMIT and IBIT at different stages of the differentiation process, revealed that a significantly higher number of cells was obtained when IMIT was used at all stages during the culture instead of IBIT (t-test p-values <0.0001 at days 25, 32 and 38, n=3), suggesting that IMIT could perform better than IBIT (Figure 1A) to differentiate iPSCs into HPCs. However, in these experiments, the cells didn’t go further than the orthochromatic erythroblast stage with both media.

**Figure 1:**
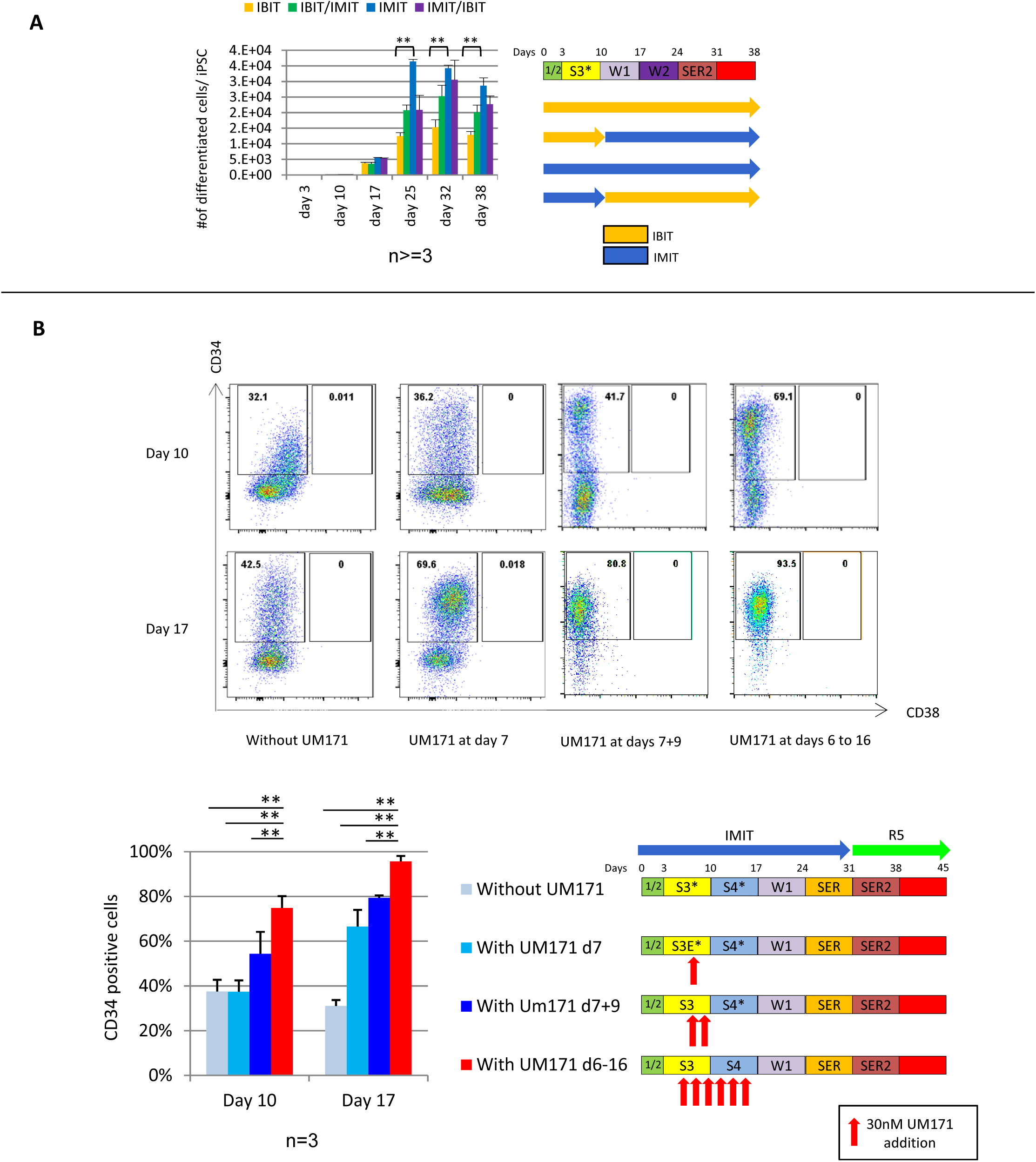
Development of IMIT and Optimization of the HSC expansion steps. **A: Right**, graph illustrates the number of cells/iPSCs observed in albumin-free IMIT or BSA-containing IBIT media during the differentiation process (n=3). **Left:** Culture conditions. 1/2 and S3: Supplement S1/S2 and S3*. The * indicates that the S3 did not include UM171 (see Table S1). Culture periods symbolized by boxes without text did not include any cytokines. The highest yield of cells was obtained when the IMIT media was used at all stages of differentiation. **B: S4 supplement.** Top: Dotplots illustrating flow cytometry analysis of the expression of CD34, CD38, before and after incubation in supplement S4 with or without stimulation with UM171. Bottom right: the timing of the addition of UM171 and culture conditions are summarized in the culture schematic. The * indicates that the S3 and S4 supplements did not include UM171 (see Table S1). Bottom left: Graph illustrating the proportion of CD34+ cells in the various conditions tested (n=3). In the presence of UM171, cells retain a much more undifferentiated phenotype, as determined by the proportion of cells that are CD34^+^CD38^−^ at day 17. The proportion of CD34+CD38− at day 17 significantly increases with increasing number of addition of UM171. When UM171 was added starting at day 6 in S3 and incorporated in the S4 supplement (at every feeding), more than 95% of the cells were CD34^+^ CD38^−^ at day 17. ** indicates that the p=value was <0.01.

We then focused on the first steps of the protocol and tested whether Trolox and methyl-β-cyclodextrin were necessary in IMIT, and whether RPMI could be used instead of IMDM (RMIT medium). This revealed that Trolox was essential and that RPMI could not be substituted for IMDM at this stage, since all cells were dead by day 10 in the absence of Trolox, or in the presence of RPMI (Figure S2A). Methyl-β-cyclodextrin had a small positive effect on the yield of cells produced that did not reach statistical significance (Figure S2A). Despite the lack of significance, we kept methyl-β-cyclodextrin in the formulation to increase the solubility of hydrophobic compounds in IMIT since these features may increase the versatility of the culture medium in future applications.

### Addition of SB431512

The first steps of the differentiation process are very sensitive to minor variations in the culture conditions and crucial for the rest of the procedure since poor differentiation at this step results in a very low overall yield. It was previously reported that blocking Activin A signaling after mesoderm induction favors definitive over primitive hematopoiesis (Kennedy et al., 2012; Ng et al., 2016). To determine if SB431512, an Activin signaling inhibitor (Inman et al., 2002), could improve the early steps of our protocol, we tested if a one-time addition of this compound to supplement S3 immediately after the dissociation step would increase the yield of the method. This revealed that the number of cells obtained between day 0 and day 10 was about 3-fold higher in the treated than in the non-treated controls independently of whether IMIT or IBIT was used (Figure S2B).

### In vitro expansion of iPSC-derived HPC

To improve on this protocol, we hypothesized that the HPCs produced in IMIT would become more developmentally mature, if they could be expanded without losing markers CD34, CD90 and CD49f. To test this hypothesis, we attempted to modify S3 by addition of either SR1 (Boitano et al., 2010) or UM171 (Fares et al., 2014), two small molecules which allow self-renewal of cord blood HSCs, and designed S4, a supplement intended to favor stem and progenitor cell expansion without differentiation. The initial formulation of S4 contained 7 cytokines, IsoButyl Methyl Xanthine (IBMX), and optionally either SR1 or UM171, (Table S1). Expansion of day-10 HPCs produced with or without adding SR1 in S3 at day 7, 9, or after day 10 in the S4 supplement were not successful because in IMIT, SR1 was relatively toxic and the HPCs could barely be maintained for 5 additional days in hematopoietic conditions (not shown). By contrast, adding a single pulse of UM171 at day 7 in S3 was not toxic and significantly increased the yield of erythroid cells (t-test p-value = 0.008, n=2, Figure S3A).

Importantly, at day 17, after 7 days of culture in supplement S4, the percentage of CD34+ cells in the UM171 culture was 66.5% ± 7% (S.D.), significantly higher than the 23.4 ± 4% obtained without exposure to UM171 (p-value at day 17 = 0.0004, n=3, Figure 1B), and the majority of the CD34+ cells had retained CD90+ and CD49f+ expression (Figure S3B) suggesting that UM171 can help expand iPSC-derived HPCs in an undifferentiated state.

Additional testing demonstrated that adding UM171 at day 7 and 9 in S3, or between days 6 and 16 in S3 and S4 had even more significant effects on the percentage of cells that retained expression of CD34+, since in the latter case CD34 expression at day 17 averaged 95± 2.4% as compared to 31.5 ± 2.3 in the absence of UM171 (p=value <0.0001 at days 10 and 17, n=3, Figure 1B). Differentiation of the cells obtained after growth in supplement S4 suggested that the cells obtained had retained their erythroid differentiation potential (see below). The protocol integrating the S4 expansion step will thereafter be referred to as the long version of the protocol to differentiate it from the shorter version without the S4 supplement.

### Erythroid differentiation of iPSC-derived HPCs

Having improved the hematopoietic specification step of the iPSC differentiation protocol, we then focused on optimizing the differentiation of iPSC-derived HPCs into cRBCs by testing different cytokine cocktails and by comparing several cell culture media.

We first compared our published conditions, culturing the HPCs in HSA containing Stemline II in the presence of the week1/week 2 cytokine cocktails (Table S1), to culturing the HPCs in the SED cocktail, and its derivative, the SER cocktail (SCF, EPO, RU486) in the IMIT and R5 media.

Initial experiments showed that the yield of cells was significantly higher (p-value <0.05 at day 25 through 38, n=2) when day 10 iPSC-derived HPCs were cultured in IMIT plus SED (condition 2) between day 10 and day 17 as compared to IMIT plus the week 1 cocktails (condition 1) (Figure 2A) and therefore that the additional cytokines present in the week 1 cocktail were not necessary. Further experiments revealed that growing day 10 iPSC-derived HPCs in condition 4 (IMIT/SED (day 10 to 17), IMIT/SER (day 17-24), R5/SER2 (day 24/31) and R5/R (day 31-38)) yielded significantly more cells than growing the cells in either condition 2 (p-value = 0.0006 at day 32, n=2) or in condition 3 (p-value = 0.009 at day 32, n=2) (Figure 2A). Thus, as described in the accompanying paper for adult erythroid culture, iPSC-derived erythroid progenitors expanded in the presence of SCF, EPO and a corticosteroid, proliferated optimally when the corticosteroid was removed before EPO and SCF (as shown when comparing condition 4 with condition 3).

**Figure 2:**
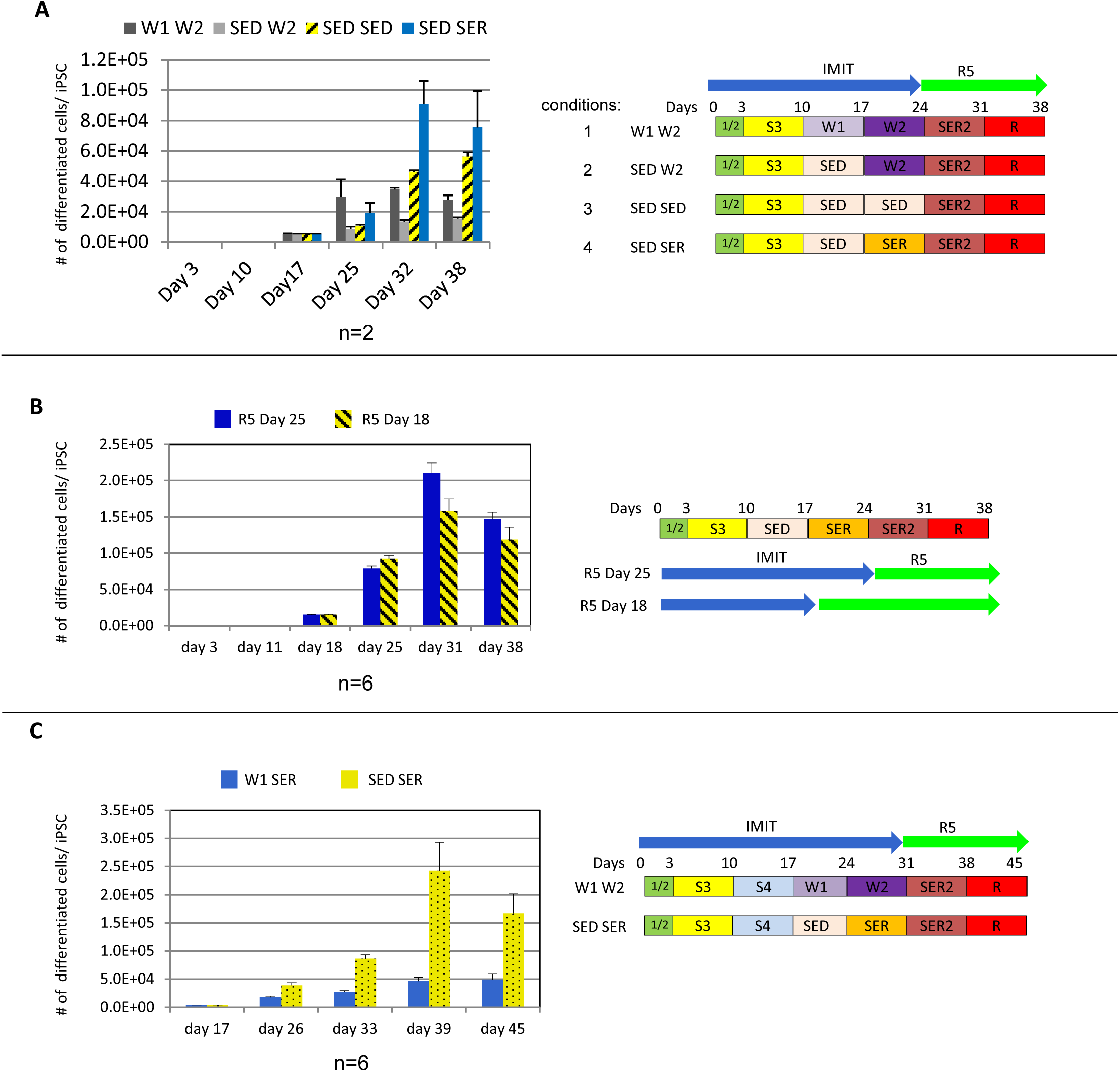
Optimization of the erythroid expansion and maturation steps (day 10 to 38) **A: Left**, diagram illustrating the number of cRBCs/iPSC produced with different cytokine combinations in order to determine the most efficient formulation and the appropriate timing to switch from glucocorticoid stimulation to glucocorticoid inhibition, n≥2. **Right:** Culture conditions schematic (short protocol). **B**: left, diagram illustrating the number of cRBCs/iPSC produced in function of the expansion medium timing switch showing that switching from IMIT to R5 at day 25 when using the SED/SER cytokines combination yields the most cells, n≥4. **Right:** Culture conditions schematic (short protocol) **C**: **Left**, diagram illustrating the number of cells/iPSC produced using the SED/SER cytokines combination in the long version of the protocol as compared with incubation in week 1/week 2 from day 17 to 31, n≥2. **Right**: culture conditions schematic (long protocol). As in A and B, erythroid differentiation for successive weeks with cytokine cocktails SED, SER, SER2 and R was most efficient.

Additional experiments confirmed that using the IMIT medium until day 24 yielded significantly more cells than switching to the R5 media at day 17 (p-value at day 31 = 0.0058, n=6, Figure 2B), and that HPCs expanded for an additional week in supplement S4 according to the long protocol also yielded significantly more cells when they were expanded in condition 4 rather than using our previous protocol (p-values = 0.043 at day 33 and <0.0001 at days 39 and 45, n=6) (Figure 2C).

### Use of small amounts of recombinant transferrin

Based on the catalog price of our media components, holo-transferrin represented more than 80% of the IMIT cost and more than 90% of the R5 cost. To decrease cost and eliminate a human-derived component, we tested whether holo-transferrin could be replaced by the cheaper commercially available recombinant human transferrin named Optiferrin.

Experiments revealed that 200 μg/mL of Optiferrin (pre-loaded with iron, or supplemented with FeIII-EDTA, see accompanying manuscript) could replace 200μg/mL of holo-transferrin (Figure 3A-B) since there were no significant differences between the yield of cells in both conditions. Reducing the Optiferrin concentration 10-fold to 20μg/mL decreased the yield by about a third (p-value = 0.06, n=2), which is an acceptable trade-off since it reduces the cost of the medium by a factor close to 10 (Figure 3A). Further experiments showed that using 50 μg/mL of Optiferrin with 4μM Fe3-EDTA in IMIT was not significantly different than 200μg/mL (not shown) while decreasing the cost of the medium by 70% (not shown).

**Figure 3:**
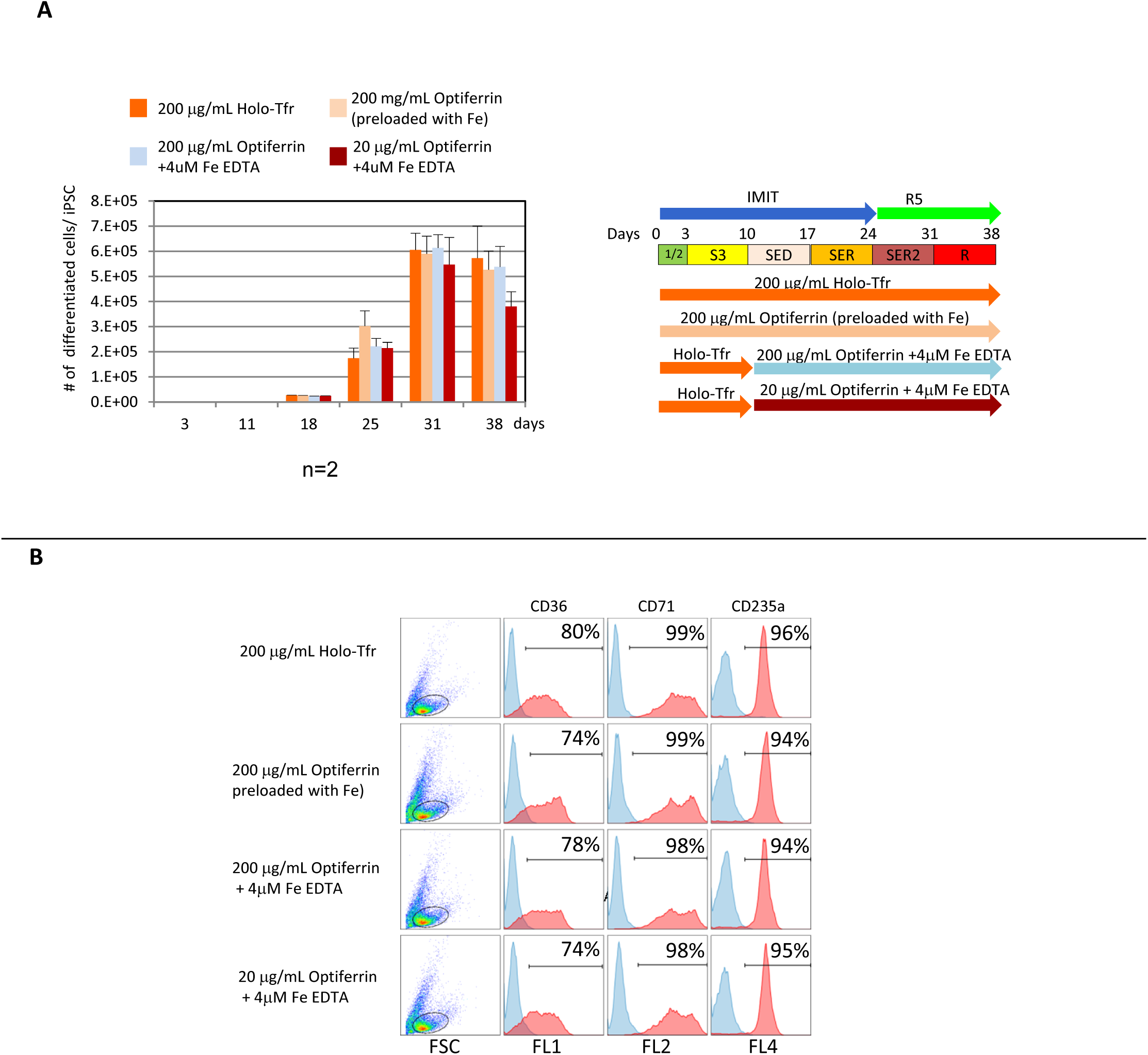
Substitution of animal derived transferrin by recombinant transferrin (Optiferrin) **A: Left:** Diagram illustrating the number of cells/iPSC observed after substituting human serum-derived holo-transferrin with Optiferrin. 200μg/ml Optiferrin (pre-loaded or supplemented with iron) can replace 200μg/mL of holo-transferrin. 20μg/ml of Optiferrin plus 4μM FeIII-EDTA can also replace holo-transferrin but with a small decrease in yield, n=2. **Right:** Culture conditions schematic. **B: FACS Analysis**. Erythroblasts obtained with transferrin or Optiferrin were analyzed by FACS at day 31. The use of Optiferrin has no effect on the immunophenotype of the cRBCs since expression of CD36, CD71 and CD235a is similar in all cases.

To determine if the source of iron and transferrin used affected differentiation, we analyzed the cells by FACS at day 31. This revealed that cells produced with holo-transferrin or Optiferrin were almost all erythroid and had a phenotypic profile very similar to that of our previously published results (Olivier et al, 2016), (Figure 3B) suggesting that the source of iron and transferrin had no major effect on differentiation. We concluded that Optiferrin can be used throughout the differentiation protocol and that medium R6 (which is medium R5 in which holo-Tf is replaced with Optiferrin and Fe3-EDTA (see Table S1) is optimal for the final differentiation steps of iPSC-derived erythroblasts.

### Robust Erythroid Differentiation (RED) Protocols

Combining these experiments, we formalized albumin-free, chemically-defined protocols that we termed short and long Pluripotent Stem Cell Robust Erythroid Differentiation (PSC-RED) protocols (Figure 4A) and characterized the cRBCs obtained. Similarly to our previous protocols, the cells obtained using the short version of the PSC-RED protocol were almost 100% orthochromatic erythroblasts but enucleated very poorly (Figure S4 and see below). Importantly, the week of expansion in S4 in the long version of the protocol resulted in the production of cells that exhibited rates of enucleation reaching more than 50% in some experiments (Figure 4B and 4C) and that averaged 41.8 ± 1.1% (n=10 independent experiments), as compared to <5% of enucleation for the short protocol. Rates of enucleation greater than 75% (n=2) (Figure 4C) could be obtained by adding a second week of culture in S4 but at the expense of a reduction in the yield of cRBCs (24000+/−3000 cRBCs/iPSC) that the higher enucleation rate was not compensating for, especially if the supplementary week of culture was taken in account.

**Figure 4:**
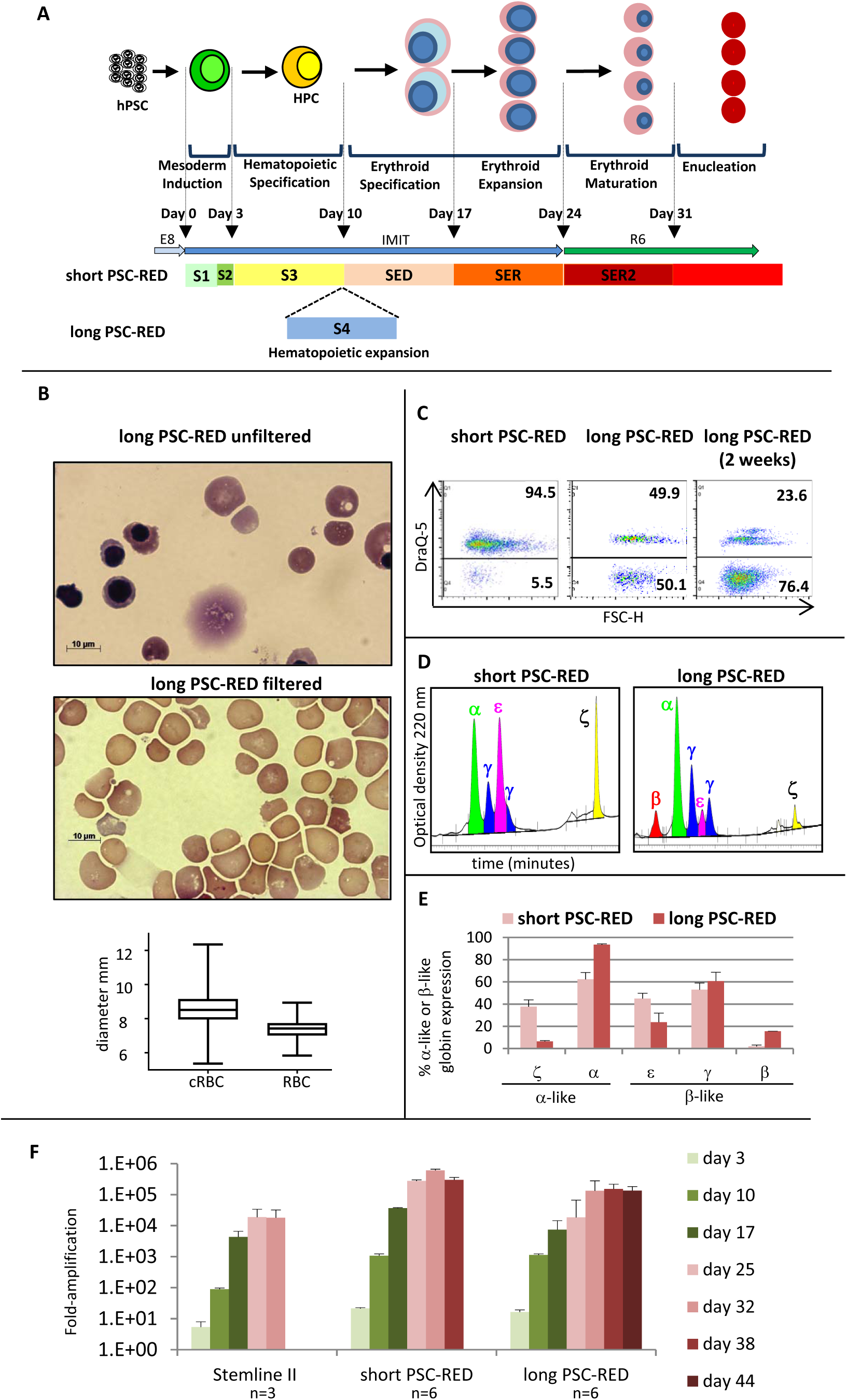
Summary of PSC-RED protocols The long PSC-RED protocol generates enucleated cRBCs from iPSCs. **A**: Graph illustrating short and long PSC-RED protocols to produce enucleated cRBCs from iPSCs. Media and supplement composition can be found in Table S1. **B**: Micrographs of rapid Ramanovsky stains illustrating the morphology of iPSC-derived enucleated cRBCs before and after filtration through a PALL Acrodisc WBC filter to eliminate the nucleated cells and the expelled nuclei. Box-and-whiskers plots on the bottom illustrate the diameter of the iPSC-derived cRBCs as compared to in vivo produced adult RBCs. The boxes represent the 25 and 75% percentile; the line inside the box the mean and the whiskers the extremes of the distribution. **C**: FACS plots illustrating enucleation rates as determined by DNA content measurement observed with the short and long PSC-RED protocols and an extra-long protocol during which the cells were incubated for two weeks in the S4 supplement. Rates of enucleation increase dramatically when iPSC-derived hematopoietic progenitors are allowed to proliferate in an undifferentiated state in the S4 supplement. **D and E**: Histograms and HPLC chromatograms illustrating globin expression of cRBCs obtained using the short or long PSC-RED protocols. Cells obtained after the long PSC-RED protocol are more developmentally mature (see also figure S4B). Globin expression phenotype is represented as percent α-like or β-like globin. Percent α-like = 100*(α or ξ)/(α+ξ). Percent β-like =100*(ε, (Gα+Aγ) or β)/(ε+Gγ+Aγ+β). Data are expressed as average ± SEM of 2 independent experiments. O.D. =optical density. **F**. Histograms illustrating the number of cRBCs/iPSC observed during differentiation in albumin containing conditions (Stemline II and IBIT) or using the short or long PSC-RED protocols. Data are expressed as average ± SEM of 3 to 6 independent experiments. Both PSC-RED protocols yield significantly more cRBCs than the albumin-containing conditions.

These important observations resolve one of the major impediments to the production of cRBCs by differentiation of iPSCs, since rates of enucleation greater than 40% are sufficient to produce pure populations of enucleated cRBC at an industrial scale. Indeed, separation by filtration of the enucleated from the nucleated cells produced using the long PSC-RED protocol (1 week in S4) yielded a 99% pure population of enucleated cells (Figure 4B).

Estimation of the size of enucleated iPSC-derived cRBCs revealed that they were about 15% larger than adult RBCs produced *in vivo* (Figure 4B) either because they were produced in culture, or because they have a more fetal phenotype.

HPLC analysis of the cRBCs produced with the short and long PSC-RED protocols revealed major differences since the short protocol yielded cells that expressed a mixture of embryonic and fetal hemoglobin (Figure 4D-E) while the long protocol yielded cells that expressed mostly γ-globin and 15.5 ± 0.16% (average ± SD) β-globin. Cells produced with the long PSC-RED protocol were therefore more developmentally mature than the cells produced with the short version of the protocol. HPLC analysis of additional experiments in which day-10 HPCs were grown in S4 for 0, 3 or 7 days, and then expanded in erythroid conditions, supported these results since expression of embryonic globins decreased and fetal and adult globin progressively increased with increasing time in HSC expansion conditions (Figure S5) independently of the cytokine cocktails used during erythroid differentiation (Figure 5E and S5).

FACS analysis of the cells obtained throughout the differentiation with antibodies against CD45, CD43, CD34, CD36, CD71 and CD235a confirmed the fundamentally different nature of the cells obtained with the short and long protocols (Figure S6). Expression of CD235a during the short protocol is detected at day 10 in most cells and continues uninterrupted until the end of the differentiation, reflecting expression of this marker in CD34+ CD43+ primitive HPCs (Vodyanik et al., 2006). This developmentally primitive phenotype is also illustrated by the lack of expression of CD45 in accordance with previous reports that demonstrated that the first wave of hematopoietic cells produced during pluripotent stem cell differentiation is CD45 negative (Qiu et al., 2005). By contrast, in the long protocol, CD235a expression almost completely disappears at day 17, reflecting the generation in supplement S4 of more definitive CD34+ CD235a-HPCs, which differentiate into CD45 positive cells by day 24 and into erythroid precursors that re-express CD235a starting at day 31. Very high levels of expression of CD36, CD71 and CD235a at the end of the differentiation confirmed that both protocols produce virtually pure populations of erythroid cells.

In terms of cell numbers, the short PSC-RED protocol generated significantly more nucleated cRBCs than our previously published procedure (p-value <0.00001, n > = 3) since it yielded up to 1 million cRBCs/iPSC in the best experiment, and an average (± S.D.) of 299,900 ± 72,444 cRBCs/iPSC as compared to 18,091 ± 24,750 for the StemLine II protocol (Figure 4F). Since there are about 5 million iPSCs per well of a six-well plate, this protocol could theoretically generate more than 10^12^ cells from a single well of iPSCs.

The long PSC-RED protocol was slightly less efficient but nevertheless yielded 135,131 ± 47,147 nucleated cRBCs/iPSC at day 45, which is significantly more than the StemLine II protocol (p-value = 0.00007, n > = 3) (Figure 4F). On average, the long PSC-RED protocol yielded about 60,000 enucleated cRBCs/iPSC at day 45. Testing of the long PSC-RED protocol on 4 additional iPSC lines that had been derived from the peripheral blood cells of four different individuals yielded similar numbers of cells and similar rates of enucleation (Figure S7).

## Discussion

We have developed chemically defined, albumin-free RED protocols that allow for the production of large numbers of enucleated cRBCs from iPSCs. The RED protocols should have an increased reproducibility compared to previously published methods because the variability associated with the unpredictable quality of animal or human-derived components is eliminated. Moreover, the media developed are easily amenable to clinical standards and should facilitate the translation process for biomedical applications.

We only observed high rates of enucleation of iPSC-derived cRBCs with the long version of the PSC-RED protocol. Similarly high rates of enucleation of RBCs derived from pluripotent stem cells have been published before (Lapillonne et al., 2010) but never with a production yield compatible with biomedical applications. While the method described here is a considerable progress in the production of enucleated cRBCs from iPSCs, it is noteworthy that elimination of extruded nuclei may be an issue that needs to be addressed for large scale production as they can clog the purification filters traditionally used in blood banking.

We have previously shown that longer periods of iPSC differentiation are associated with the production of fetal hemoglobin rather than embryonic hemoglobin and therefore that lengthening iPSC differentiation time recapitulated some aspects of normal erythroid development (Qiu et al., 2008). HPLC analysis of globin expression of cRBCs obtained with the short and long PSC-RED protocols clearly show that a week expansion of HPCs in S4 is sufficient to obtain more developmentally mature cRBCs and suggests that the increase in the rate of enucleation reflects the emergence or the preferential amplification of more definitive progenitors during the week of culture in S4. The increased overall length of the protocol might also select for more developmentally mature cells, since primitive cells likely differentiate and die before the end of the culture. Finally, elimination of inhibitory compound(s) present in undefined media components might also contribute to the increased rate of enucleation.

The long PSC-RED protocol will be particularly useful to generate drug-carrying cRBCs since it starts from iPSCs, an immortal cell type that is highly amenable to precise genetic modifications. As discussed in the introduction, clinical applications of the technology based on cRBCs genetically engineered to express therapeutic molecules, are particularly attractive, because they require much smaller numbers of cells than transfusion applications.

The cost of cRBC production for clinical applications is a major limitation of the technology. Since culture volumes increase exponentially during cRBC production, production costs are mostly incurred during the last weeks of culture. Importantly, the last four weeks of the PSC-RED protocol require only a commonly used base media, a few small molecules, low amounts of recombinant transferrin, which is cheaper than human plasma-derived transferrin, and two cytokines, EPO and SCF, which are in the public domain and relatively simple to produce in large amounts. In addition to yielding enucleated cells, the PSC-RED protocol is therefore relatively inexpensive. This low cost will help make the many applications of high-value genetically modified cRBCs economically feasible.

