## Supplementary Figures and Tables Legends for "PSC-RED, an Albumin-Free Robust Erythroid Differentiation Method to Produce Enucleated Red Blood Cells from Human Pluripotent Stem Cells"

**Supplementary Figure Legends:**

**Figure S1:** Phase contrast micrographs (x40 magnification) of cells in culture during the first 10 days of differentiation.

At day 0 the cells are small colonies of iPSC undifferentiated, by day 2 they already exhibit a changed morphology with larger and more elongated cells. The differentiation continues on day 3 with the emergence of endothelial looking cells. On day 4 after dissociation and replating the cells arrange in small clusters of elongated cells, from these cluster on days 5, 6 and 7, emerge round and bright cells. The round and bright cells divide very actively. Tens of millions of cells can be obtained by day 10 when starting with 100,000 iPSCs in 1 well at day 0.

**Figure S2: Development of IMIT**

**A:** Components of the IMIT medium were tested for their importance in supporting iPSC differentiation toward erythroid cells. Top: Schematic of the differentiation conditions. Bottom: bar graph illustrates the number of differentiated cells/iPSC obtained after 38 days of differentiation (n=2). Trolox and the use of IMDM (rather than RPMI) are absolute requirements since all cells were dead after day 10 (compare IMIT control, with RMIT or no Trolox). Methyl-β-cyclodextrin (MbC) had a noticeable but non-significant effect on the yield of cells.

**B: Left,** bar graph illustrating the increased number of cells at day 10 after a single pulse of SB431542 at day 3 whether IBIT or IMIT is used as expansion medium. **Right:** Schematic of the differentiation conditions.

**Figure S3: Optimization of the HSC expansion steps**

**A: Left** Diagram summarising the effect of a single addition of UM171 on the number of cells/iPSC produced in an early version of the long differentiation protocol, n≥2. **Right:** Culture condition schematic.

**B**: Dotplots illustrating flow cytometry analysis of the expression of CD34, CD38, CD45RA, CD90 and CD49f after a single dose of UM171 at day 7 as described in the graph above. In the presence of UM171, cells retain a much more undifferentiated phenotype, as determined by the proportion of cells expressing CD34, CD49f and CD90 at day 17, data representative of 2 or more experiments.

**Figure S4:** Micrographs of rapid Ramanovsky stain illustrating the morphology of iPSC-derived cRBCs obtained at the end of the short PSC-RED protocol. Most of the cells are orthochromatic erythroblasts and have a primitive morphology.

**Figure S5: Expansion and globin expression as a function of time in S4**

**Top:** Culture conditions schematic.

**Bottom left:** Diagram illustrating the yield of cells/iPSC for up to 45 days either without, or with 3 or 7 days incubation in supplement S4 (containing UM171). **Middle**, HPLC chromatograms, and **right** bar graph illustrating the proportion of embryonic (epsilon and zeta), fetal (gamma and alpha) and adult globin (beta and alpha) produced as a function of time in S4 supplement. Progenitors cultured the longest in S4, exhibit a more mature phenotype characterized by a sharp decrease in epsilon- and zeta-globin chain expression, and an increase in alpha-, gamma- and beta-globin chain expression, n≥4.

**Figure S6: Phenotypic analysis of the cell markers during the PSC-RED protocols**

**A: Flow cytometry analysis during the Short PSC-RED protocol:**

Representative flow cytometry plots depicting the expression of hematopoietic and erythroid markers during the short PSC-RED protocol. Cells are gated by size (FSC) and granularity (SSC) to exclude debris, dead and dying cells. For each marker the corresponding isotype control is shown in blue. At day 10, the cells are almost all hematopoietic as indicated by CD34 and CD43. During the differentiation, CD235a is expressed on most cells at all time and expression of CD45 never takes off. CD36 expression culminates at day 31 before receding during the cell maturation process. At day 38 the cells are almost all erythroid as indicated by CD36, CD71 and CD235a expression.

The culture conditions schematic is shown under the plot and the data are representative of 3 or more experiments.

**B: Flow cytometry analysis during the long PSC-RED protocol:**

Representative flow cytometry plots depicting the expression of hematopoietic and erythroid markers during the long PSC-RED protocol. Cells are gated by size (FSC) and granularity (SSC) to exclude debris, dead and dying cells. For each marker the corresponding isotype control is shown in blue. Cells are almost 100% CD45 positive at day 24 but loose expression of this marker as erythroid differentiation progresses. Starting at day 10, most cells in the culture are hematopoietic as determined by expression of CD43. Expression of CD34 peaks at day 17, when it reaches almost 100%. Expression of this marker decreases rapidly when the cells are placed in the SED conditions. CD36 expression is first detected at day 31 and decreases at day 38 as the cells enter their final maturation. CD71 expression is complex because this marker is expressed in all cells at low levels, at very high levels in erythroid cells except in orthochromatic erythroblasts and reticulocytes where expression is low again. CD235a expression is high in day 10 HPCs, low or absent at day 17 and 24 and high again at day 31 and 38. Combined expression of CD36, 71 and 235a demonstrates that almost all cells are erythroid at day 38.

The culture conditions schematic is shown under the plot and the data are representative of 3 or more experiments.

**Figure S7: Production of enucleated cRBCs from 4 different iPSC lines.**

**A:** Diagram illustrating the percentage of enucleated cells at days 43 and 45 (n=2) as measured by Draq5 staining. Cells were obtained by differentiation of 4 different iPSC lines generated from four different individuals using the long PSC-RED protocol.

B: Micrographs of rapid Ramanovsky stain illustrating the morphology of the cRBCs obtained from the four iPSC lines at the end of the long PSC-RED protocol after filtration through a Pal Acrodisc 25mm WBC filter.

Tables:

| | | | | | |
|---|---|---|---|---|---|
| **Culture media and supplements** | **IBIT**<br>IMDM with 1mM Glutamine<br>BSA 1%<br>Insulin 10µg/mL<br>Transferrin 200µg/mL<br>β-mercapto-ethanol 0.1mM<br>Lipids (1X)<br>Ethanolamine<br><br>**IMIT**<br>IMDM with 1mM Glutamine<br>Methyl-β-Cyclodextrin 0.1mg/mL<br>Trolox50µM<br>Insulin 10µg/mL<br>Optiferrin 50µg/mL<br>FeIII-EDTA 4µM<br>Lipids (1.5X)<br>Ethanolamine<br><br>**RIT**<br>RPMI 1640 with 1mM Glutamine<br>Insulin 10µg/mL<br>Transferrin 200µg/mL<br>β-mercapto-ethanol 0.1mM<br>Lipids (1/200)<br>Ethanolamine | **R5**<br>RPMI 1640<br>L-ascorbic acid 220 uM<br>Insulin 10ug/mL<br>Transferrin 200ug/mL<br>Lipids 1/200<br><br>**R6**<br>RPMI 1640<br>L-ascorbic acid 220 uM<br>Insulin 10ug/mL<br>Optiferrin 20 ug/mL<br>FeIII-EDTA 4µM<br>Lipids 1/200 | **S1**<br>BMP4 10ng/mL<br>VEGF 165 10ng/mL<br>Wnt3A/5A 5ng/mL each<br>Activin A 5ng/mL<br>Inhibitor VIII 2uM<br>bFGF 10ng/mL<br><br>**S2**<br>BMP4 20ng/ml<br>VEGF 165 30ng/mL<br>Wnt3A/5A 5ng/mL each<br>Activin A 5ng/mL<br>Inhibitor VIII 2uM<br>bFGF 10ng/mL<br>SCF 20ng/mL<br>β-Estradiol 0.4ng/mL | **S3**<br>BMP4 20ng/mL<br>VEGF 165 30ng/mL<br>bFGF 20ng/mL<br>SCF 30ng/mL<br>TPO 10ng/mL<br>IGF2 10ng/mL<br>β-Estradiol 0.4ng/mL<br>SB431542 3µM on day 3 only<br>IBMX 50 µM<br>UM171 30nM after day 6<br>Heparin 5µg/mL<br><br>**S4**<br>VEGF165 5ng/mL<br>bFGF 5ng/mL<br>SCF 15ng/mL<br>TPO 10ng/mL<br>IGF2 10ng/mL<br>IBMX 30 µM<br>PDGF AB 5ng/mL<br>ANGPTL5 5ng/mL<br>CCL28 5ng/mL<br>UM171 30nM<br>Heparin 5µg/mL<br><br>**S3*** and **S4*** in some of the figures refers to supplements S3 and S4 without UM171 that were used during the development of the PSC-RED protocols. | **SED**<br>SCF 100ng/mL<br>EPO 4U/mL<br>IBMX 50 µM<br>Dexamethasone 1 µM<br><br>**SER**<br>SCF 50ng/mL<br>EPO 4U/mL<br>RU486 1 µM<br><br>**SER2**<br>SCF 10ng/mL<br>EPO 4U/mL<br>RU486 1 µM<br><br>**R**<br>RU486 1 µM<br><br>**Week 1 (W1)**<br>StemSpan SFEM<br>Hydrocortisone 1µM<br>SCF 50 ng/mL<br>FLT3L 16.7 ng/mL<br>IL3 6.67 ng/mL<br>EPO 1.33 U/mL<br><br>**Week 2 (W2)**<br>StemSpan SFEM<br>Hydrocortisone 1µM<br>SCF 20 ng/mL<br>IGF1 20 ng/mL<br>IL3 6.67 ng/mL<br>EPO 2 U/mL<br><br>**SEII cytokines**<br>SCF 4.4 ng/mL<br>EPO 3u/mL<br>IGF1 4.4ng/mL<br>IL3 1.5 ng/mL |

**Table S1: Media and supplements**

Table S2:

| **Reagent** | **Provider** | **Catalog Number** |
|---|---|---|
| IMDM with 1mM Glutamine | Biochrom | FG0465 |
| RPMI 1640 with 1mM Glutamine | Gibco | 61870 |
| StemSpan SFEM | Stemcell Technologies | 09650 |
| Methyl- β -Cyclodextrin | Sigma | C4555 |
| Trolox | Sigma | 238813 |
| Insulin | Sigma | I9218 |
| Chemically defined Lipids 200X | Gibco | 11905 |
| Ethanolamine | Sigma | E0135 |
| BSA | Gibco | From Kit A1000701 |
| β-mercapto-ethanol 1000X | Gibco | 21985 |
| L-ascorbic acid | Sigma | A8960 |
| Holo-Transferrin | R&D Systems/Biotechne | 2914-HT |
| Optiferrin | FisherScience | NC9954311 |
| FeIII-EDTA | Sigma | E6760 |
| BMP4 | R&D Systems/Biotechne | 314-BP |
| VEGF165 | Peprotech | 100-20 |
| Wnt3A | R&D Systems/Biotechne | 5036-WN |
| Wnt5A | R&D Systems/Biotechne | 645-WN |
| Activin A | Peprotech | 120-14 |
| GSK3β Inhibitor VIII | Calbiochem/EMD Millipore | 361549 |
| aFGF | Peprotech | 100-17A |
| bFGF | Peprotech | 100-18B |
| SCF | Peprotech | 300-07 |
| β-Estradiol | Sigma | E2758 |
| TPO | Peprotech | 300-18 |
| IGF1 | Alfa Aesar | BT-106 |
| IGF2 | Alfa Aesar | BT-107 |
| SB431542 | Tocris/Biotechne | 1614 |
| UM171 | Stemcell Technologies | 72912 |
| IBMX | Sigma | I5879 |
| PDGF AB | Peprotech | 100-00AB |
| ANGPTL5 | R&D Systems/Biotechne | 6675-AN |
| CCL28 | Peprotech | 300-57 |
| Heparin | Sigma | H3149 |
| EPO | Amgen | NDC 55513-126-10 |
| Dexamethasone | Sigma | D4902 |
| RU486 | Sigma | M8046 |
| Hydrocortisone | Sigma | H0888 |
| FLT3L | Peprotech | 300-19 |
| IL3 | Peprotech | 200-03 |
| GM-CSF | Peprotech | 300-03 |
| G-CSF | Peprotech | 300-23 |

**Table S2: Reagents**

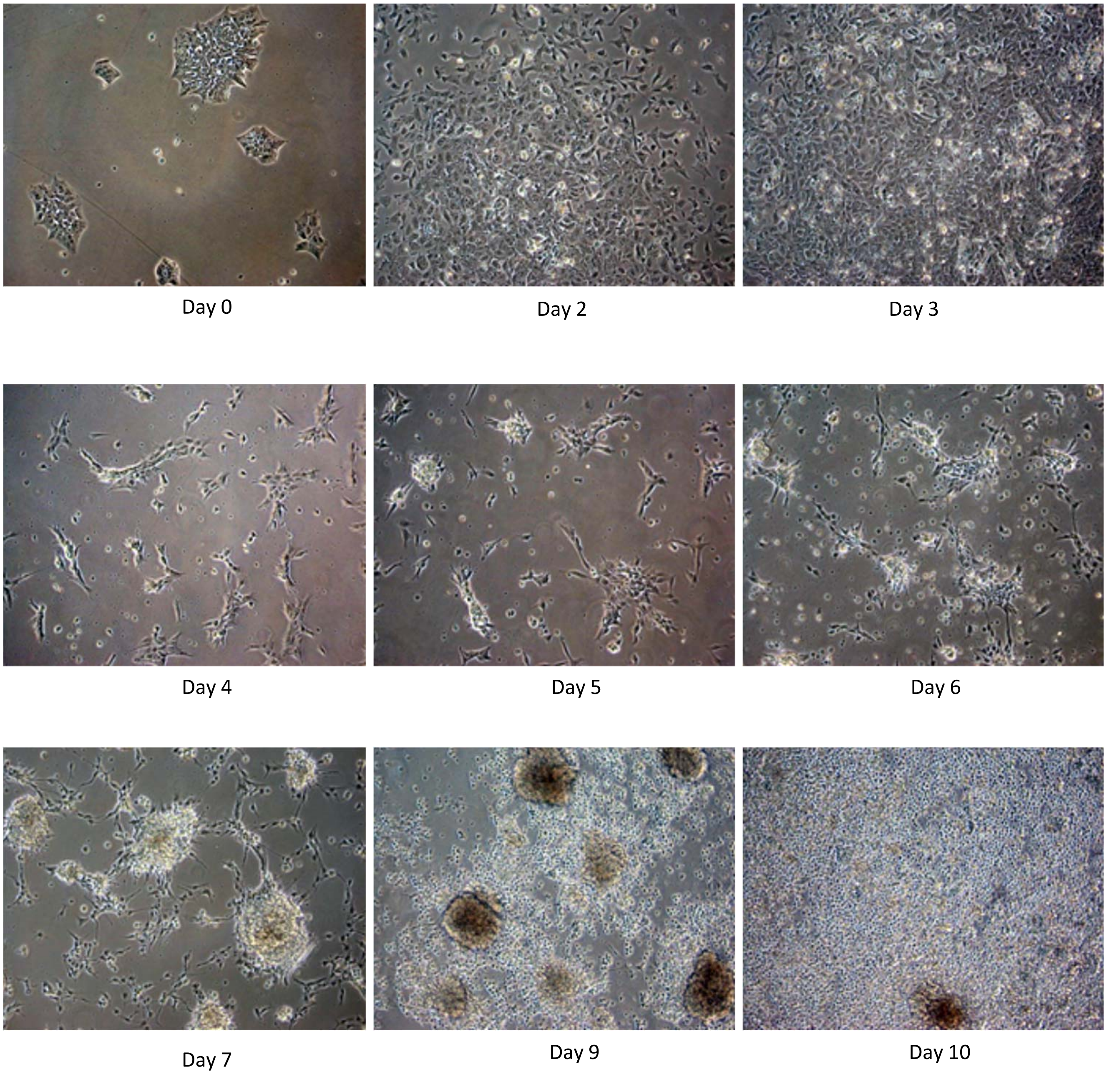


Figure S1

A

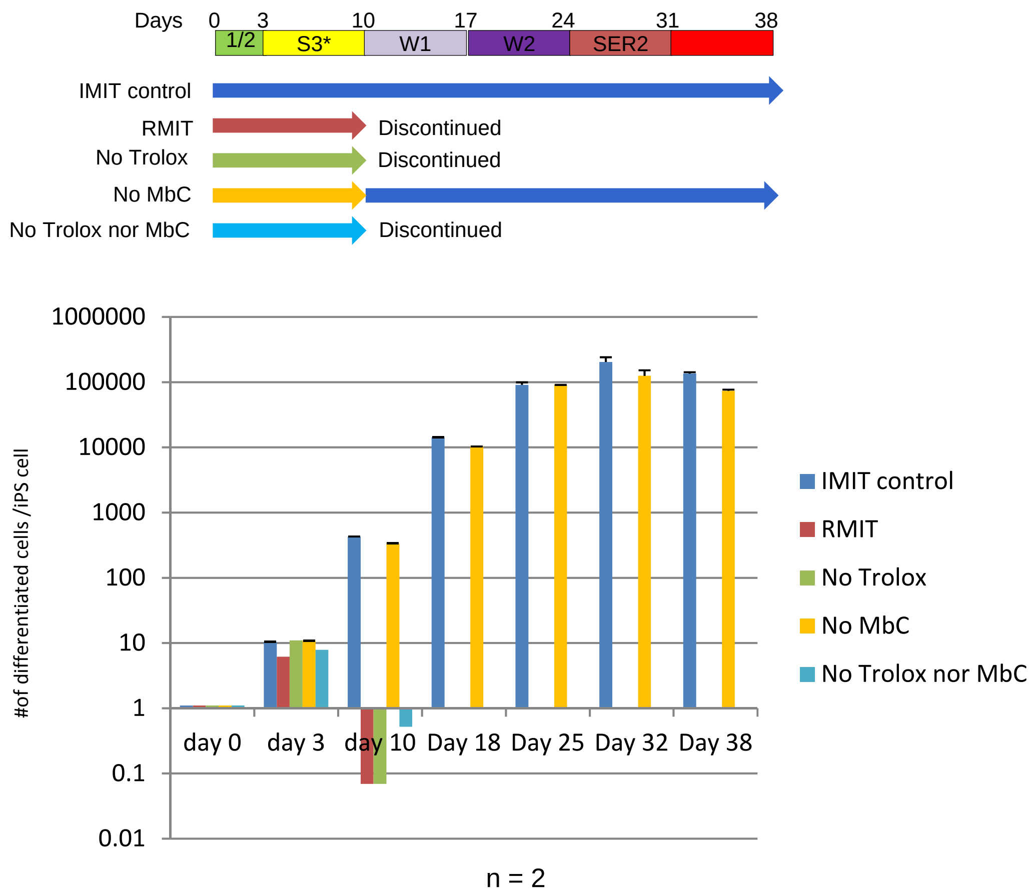
Days
0
3
10
17
24
31
38
1/2
S3*
W1
W2
SER2
IMIT control
RMIT
Discontinued
No Trolox
Discontinued
No MbC
No Trolox nor MbC
Discontinued
#of differentiated cells /iPS cell
1000000
100000
10000
1000
100
10
1
0.1
0.01
day 0
day 3
day 10
Day 18
Day 25
Day 32
Day 38
IMIT control
RMIT
No Trolox
No MbC
No Trolox nor MbC
n = 2

B

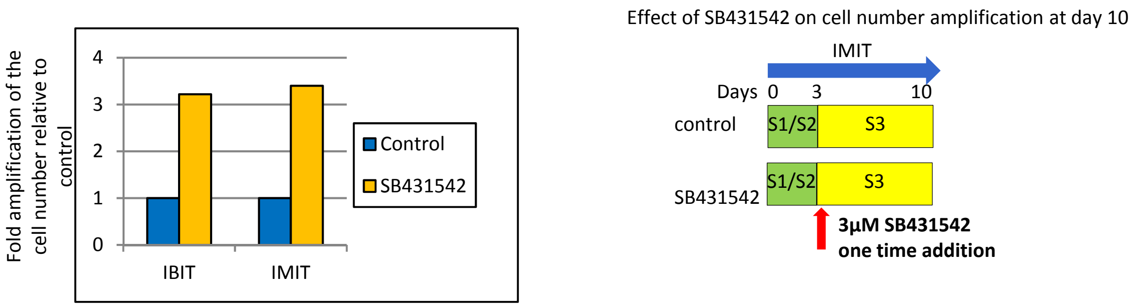
Fold amplification of the cell number relative to control
4
3
2
1
0
IBIT
IMIT
Control
SB431542
Effect of SB431542 on cell number amplification at day 10
IMIT
Days
0
3
10
control
S1/S2
S3
SB431542
S1/S2
S3
3µM SB431542
one time addition

Figure S2

A

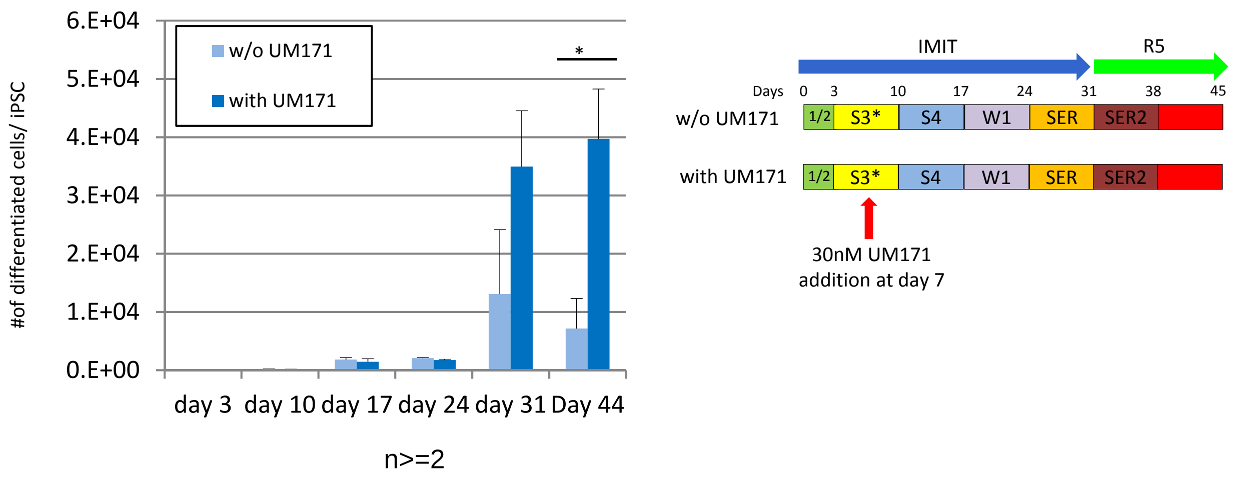
w/o UM171
with UM171
6.E+04
5.E+04
4.E+04
3.E+04
2.E+04
1.E+04
0.E+00
#of differentiated cells/ iPSC
*
day 3
day 10
day 17
day 24
day 31
Day 44
n>=2
IMIT
R5
Days
0
3
10
17
24
31
38
45
w/o UM171
with UM171
1/2
S3*
S4
W1
SER
SER2
30nM UM171
addition at day 7

B

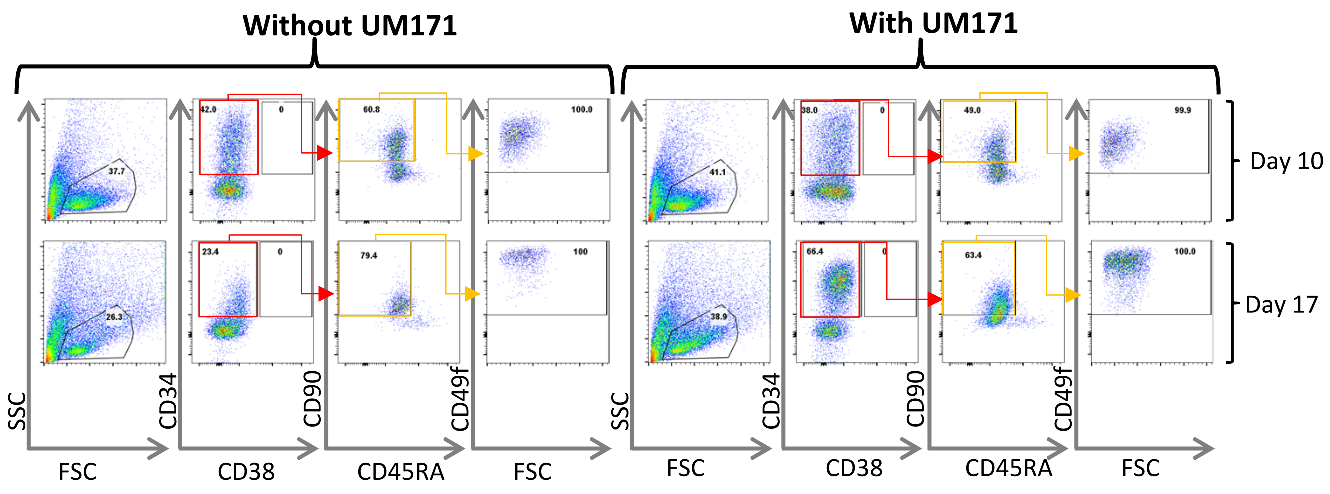
Without UM171
With UM171
Day 10
Day 17
SSC
FSC
CD34
CD38
CD90
CD45RA
CD49f
FSC

Figure S3

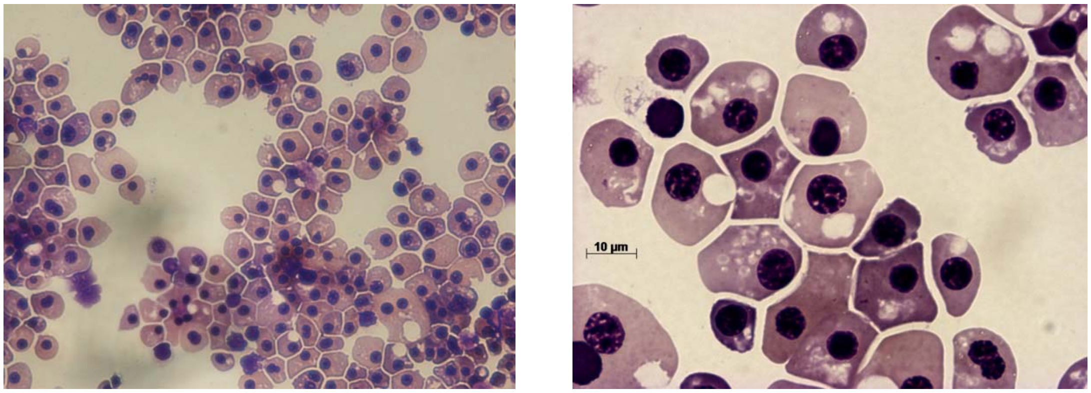


Figure S4

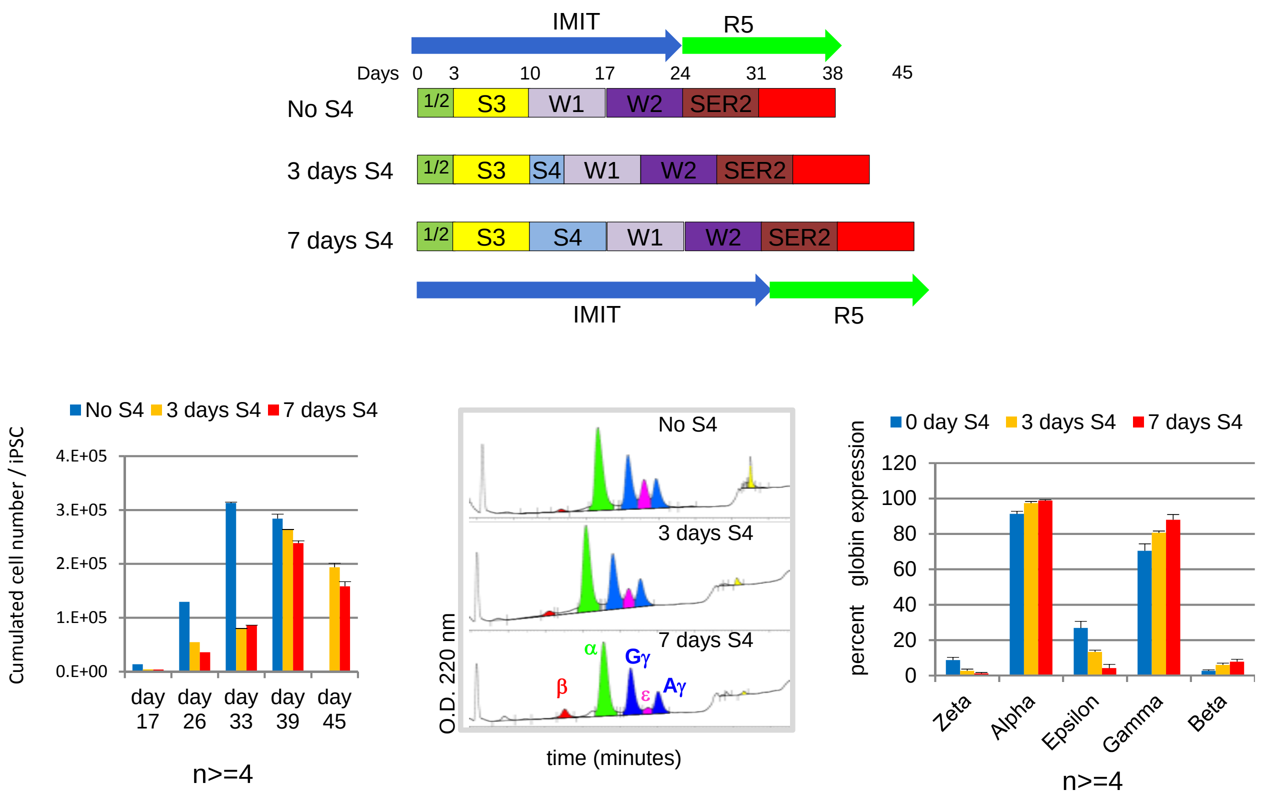
IMIT
R5
Days 0 3 10 17 24 31 38 45
No S4
1/2 S3 W1 W2 SER2
3 days S4
1/2 S3 S4 W1 W2 SER2
7 days S4
1/2 S3 S4 W1 W2 SER2
IMIT
R5
No S4 3 days S4 7 days S4
Cumulated cell number / iPSC
4.E+05
3.E+05
2.E+05
1.E+05
0.E+00
day 17
day 26
day 33
day 39
day 45
n>=4
No S4
3 days S4
7 days S4
α
Gγ
β
ε
Aγ
O.D. 220 nm
time (minutes)
0 day S4 3 days S4 7 days S4
percent globin expression
120
100
80
60
40
20
0
Zeta
Alpha
Epsilon
Gamma
Beta
n>=4

Figure S5

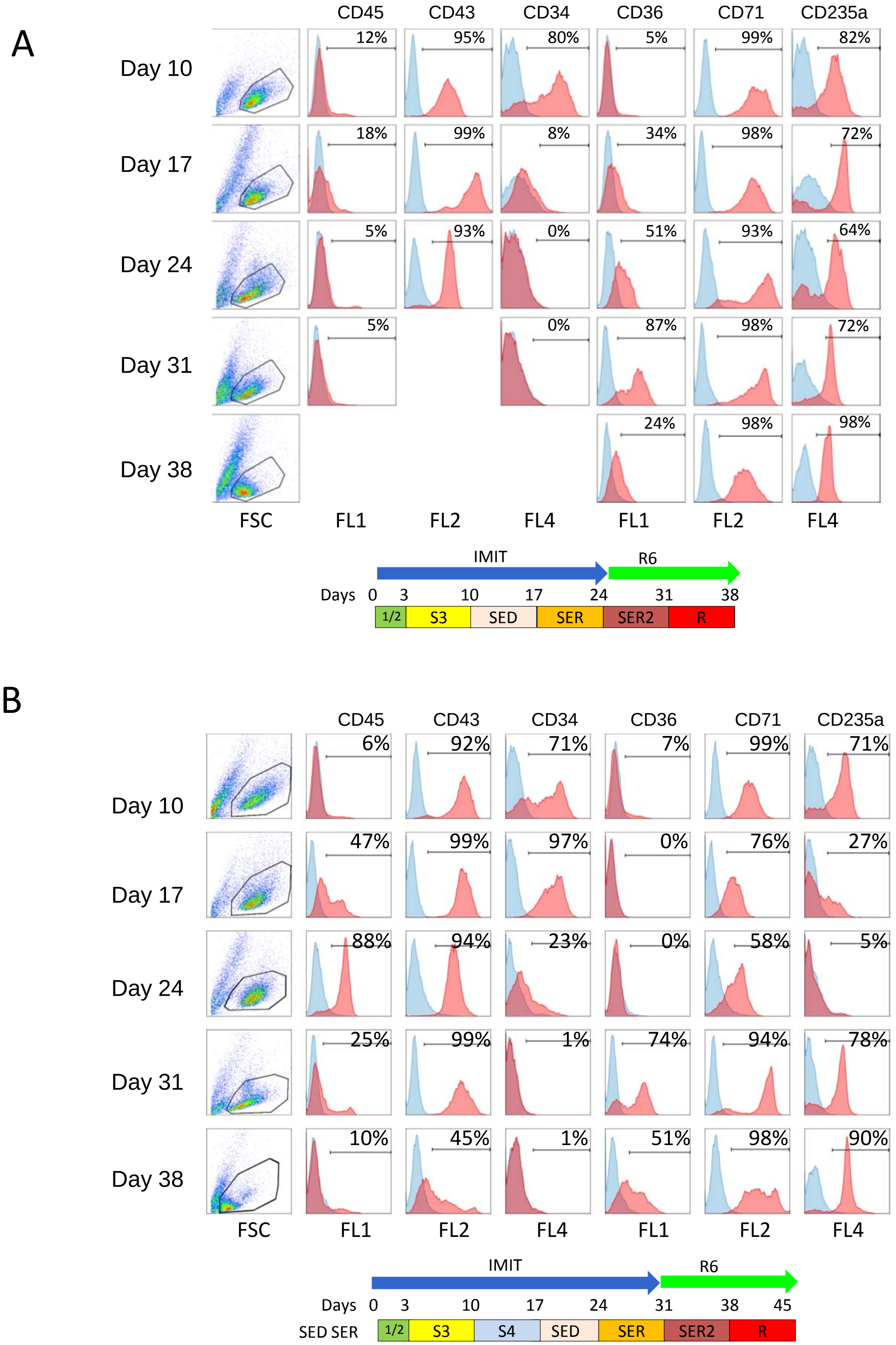
A
CD45
CD43
CD34
CD36
CD71
CD235a
Day 10
12%
95%
80%
5%
99%
82%
Day 17
18%
99%
8%
34%
98%
72%
Day 24
5%
93%
0%
51%
93%
64%
Day 31
5%
0%
87%
98%
72%
Day 38
24%
98%
98%
FSC
FL1
FL2
FL4
FL1
FL2
FL4
IMIT
R6
Days
0
3
10
17
24
31
38
1/2
S3
SED
SER
SER2
R
B
CD45
CD43
CD34
CD36
CD71
CD235a
Day 10
6%
92%
71%
7%
99%
71%
Day 17
47%
99%
97%
0%
76%
27%
Day 24
88%
94%
23%
0%
58%
5%
Day 31
25%
99%
1%
74%
94%
78%
Day 38
10%
45%
1%
51%
98%
90%
FSC
FL1
FL2
FL4
FL1
FL2
FL4
IMIT
R6
Days
0
3
10
17
24
31
38
45
SED SER
1/2
S3
S4
SED
SER
SER2
R

Figure S6

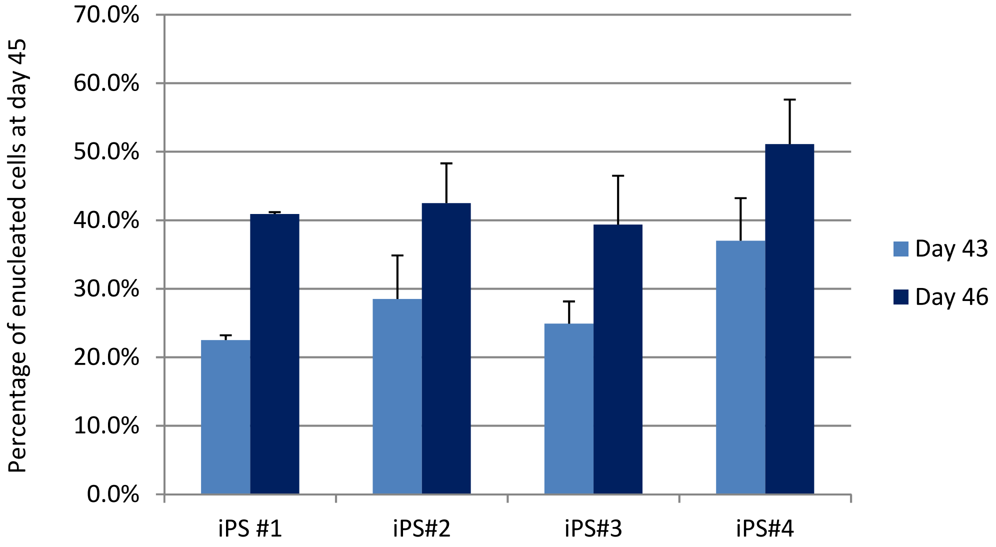


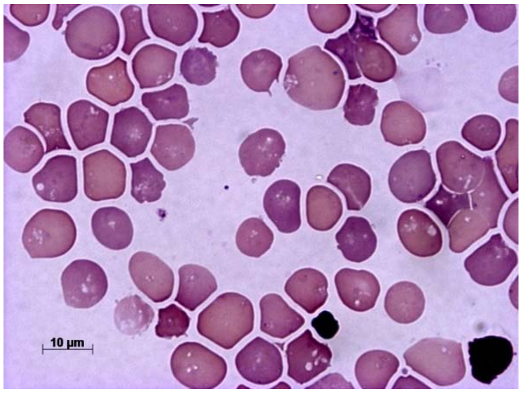


iPS#OM1

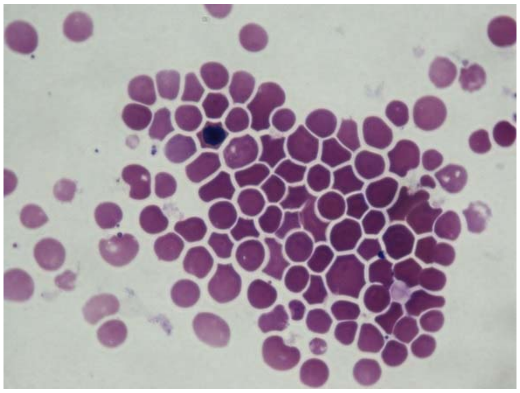

iPS#OM3

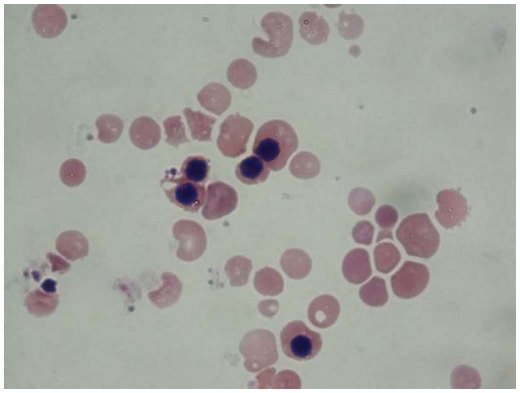

iPS#OM2

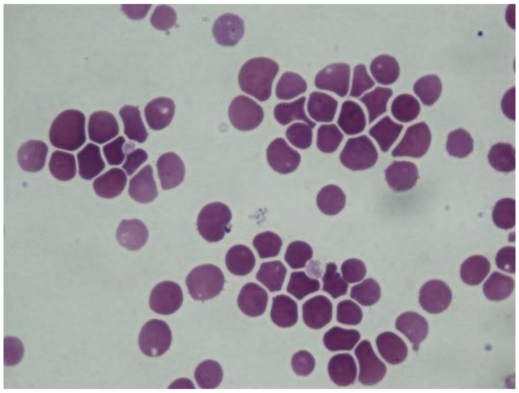

iPS#OM4

Figure S7
